## Supplemental Figures for "Location bias contributes to functionally selective responses of biased CXCR3 agonists"

### SUPPLEMENTAL FIGURE 1

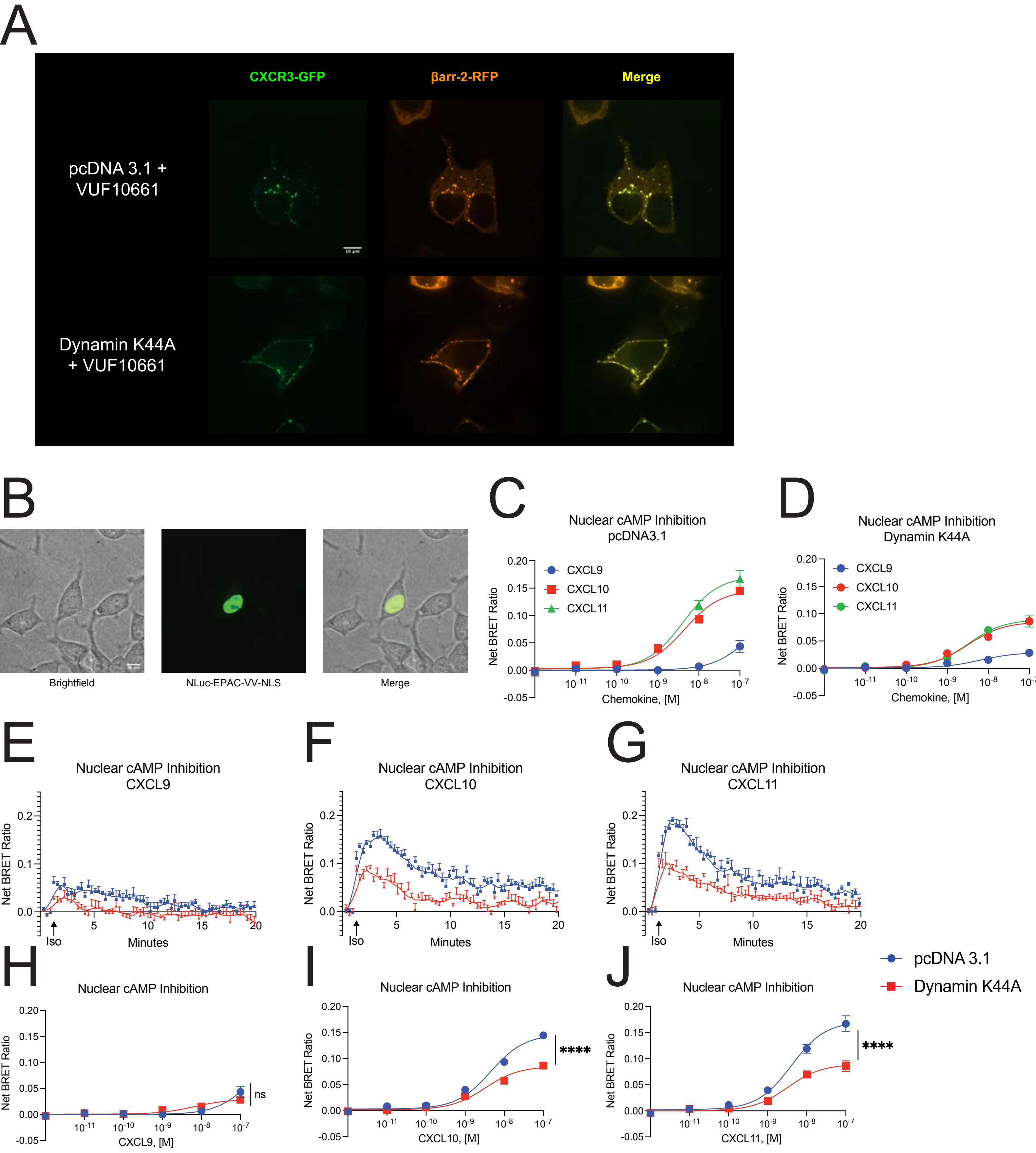

SUPPLEMENTAL FIGURE 2

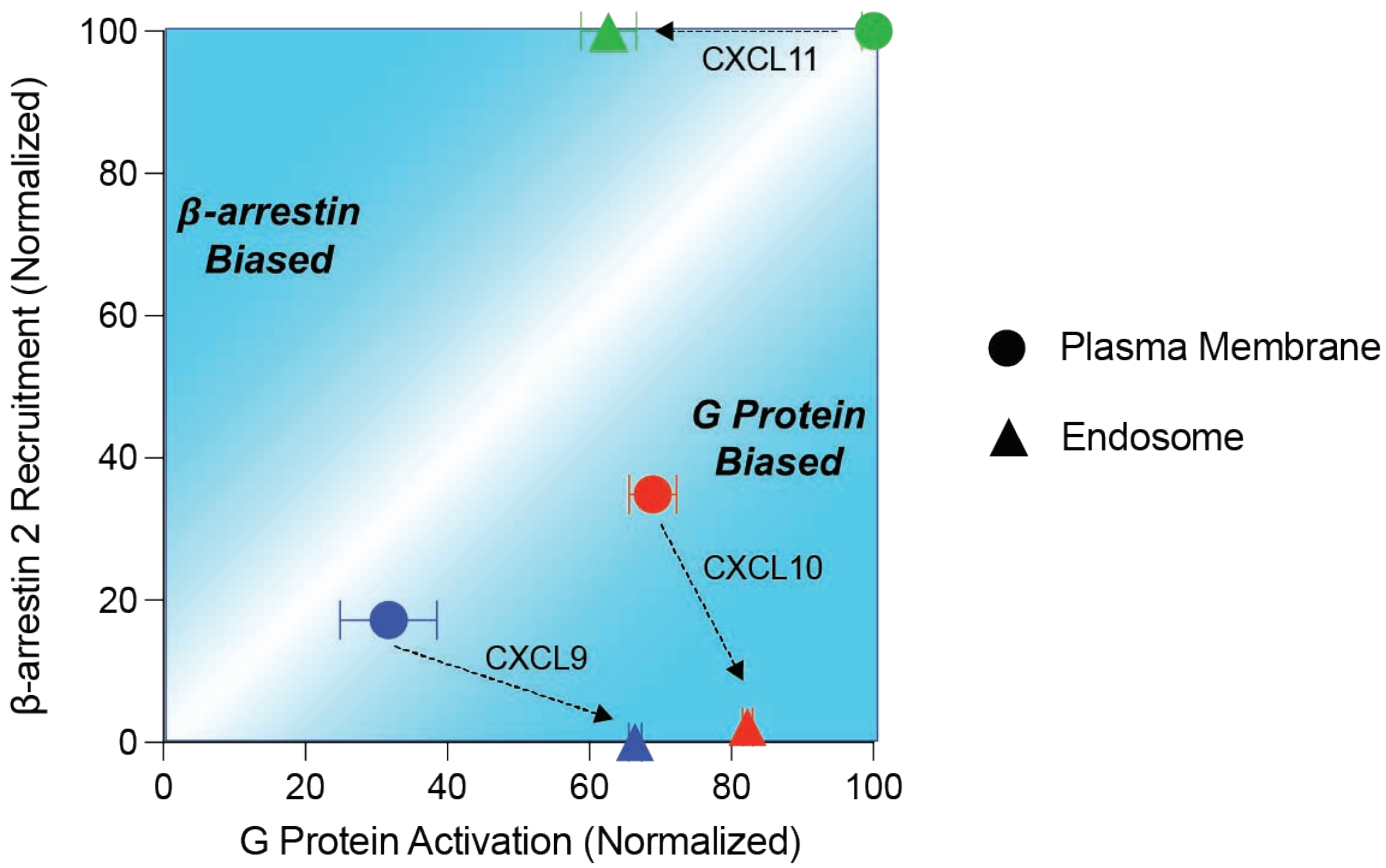

### SUPPLEMENTAL FIGURE 3

A

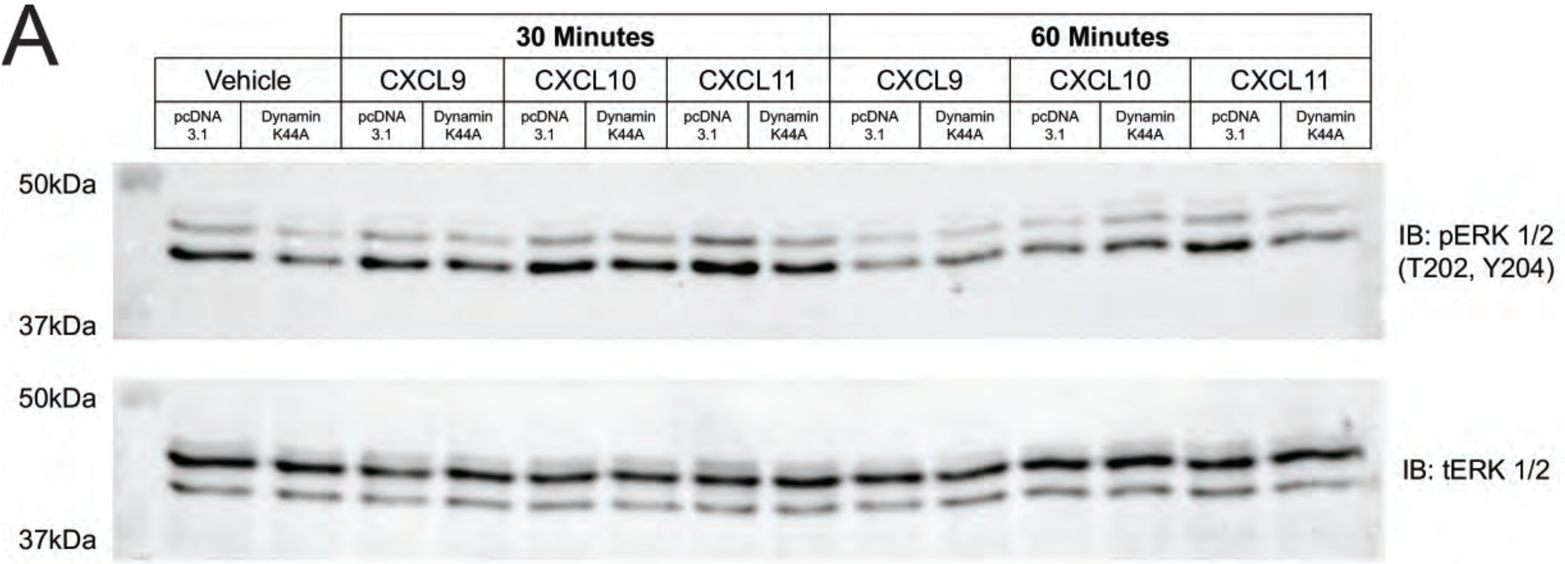


B

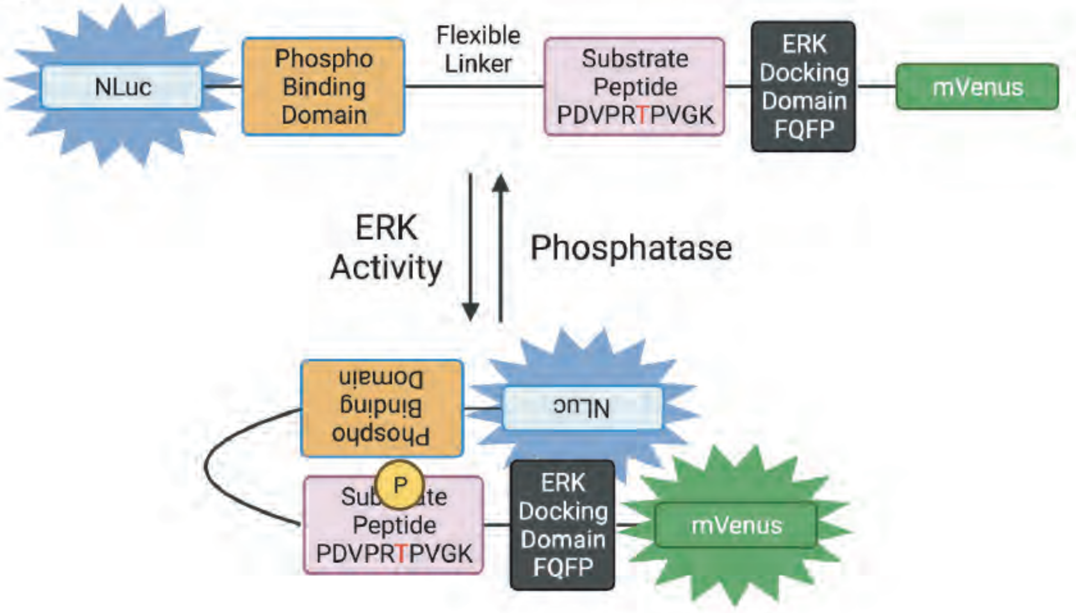


C

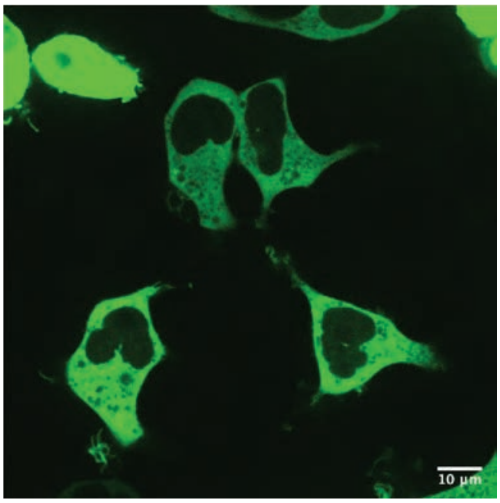


Cytosolic ERK Sensor

D

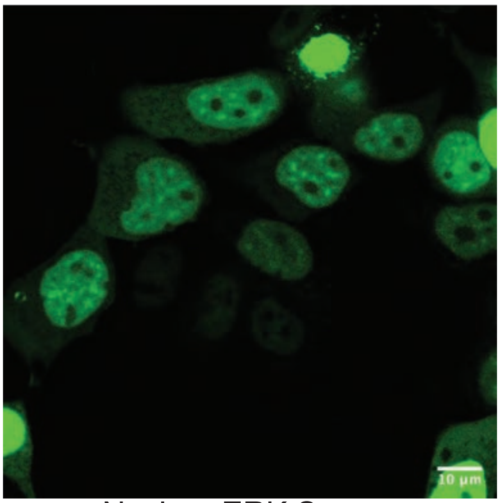


Nuclear ERK Sensor

### SUPPLEMENTAL FIGURE 4

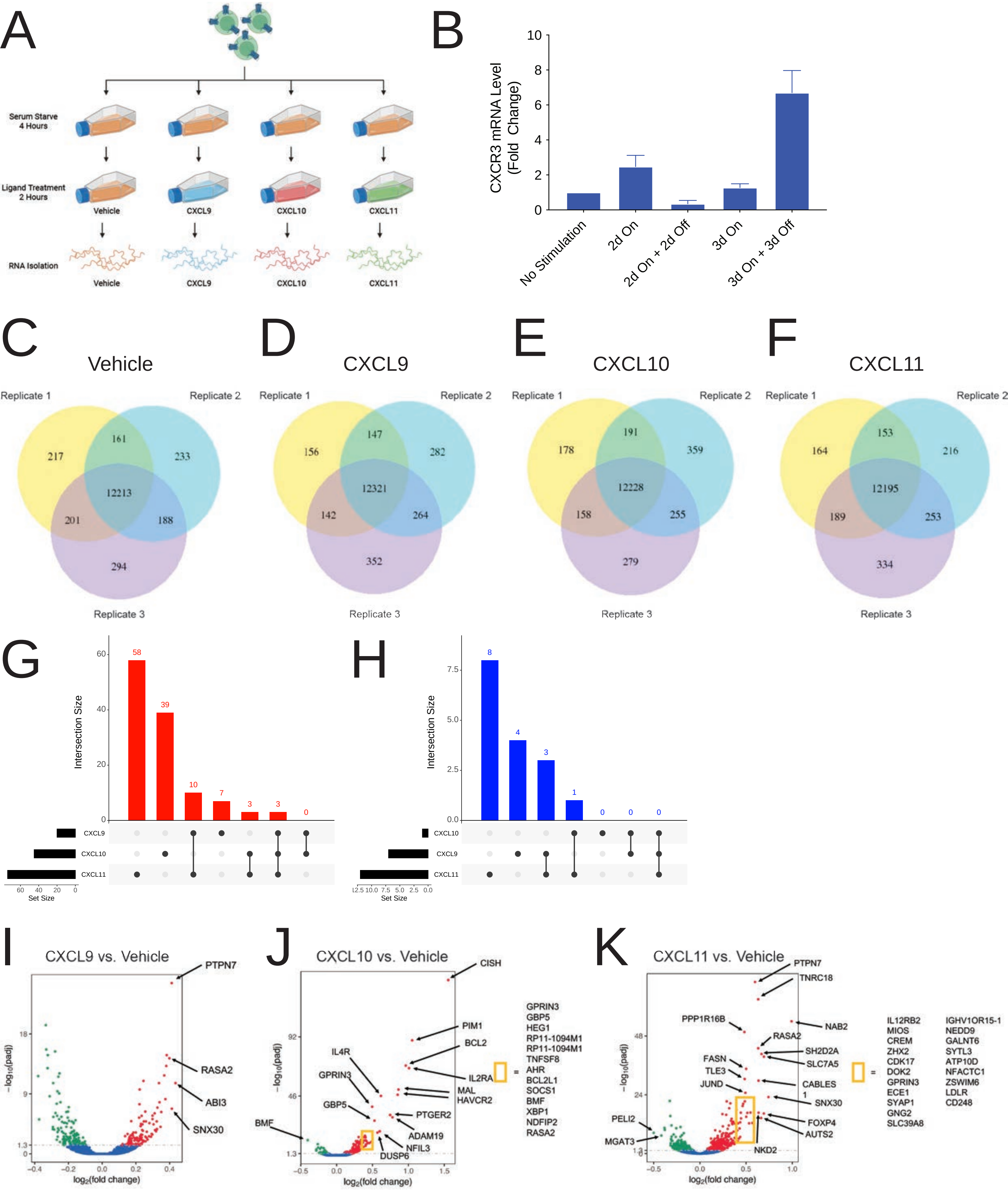

### SUPPLEMENTAL FIGURE 5

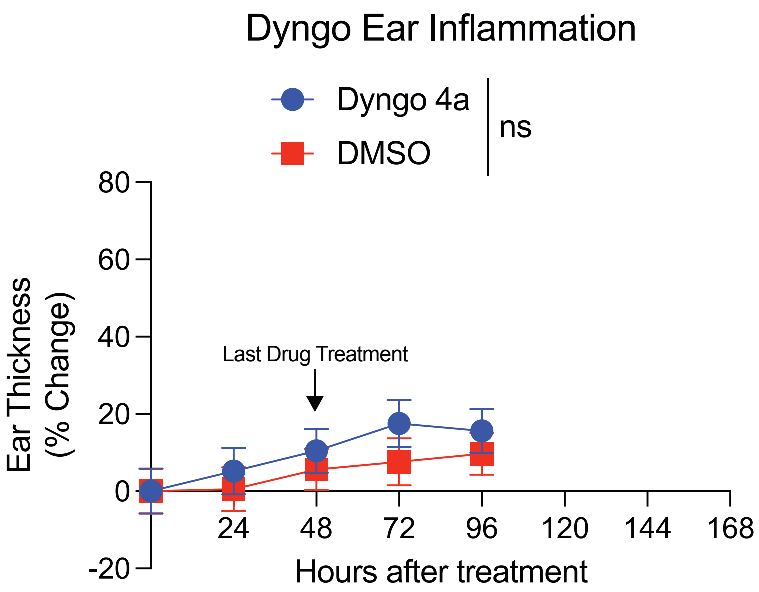
